# Endocannabinoid-dependent persistent decrease of GABAergic transmission on dopaminergic neurons underlies gene-environment interaction-induced susceptibility to cocaine sensitization

**DOI:** 10.1101/2022.08.10.503451

**Authors:** Valeria Serra, Francesco Traccis, Sonia Aroni, Marco Bortolato, Miriam Melis

## Abstract

Vulnerability to develop cocaine use disorder depends upon an unpredictable combination of genetic and non-genetic risk factors. Early life adversity and adolescence are critical non-genetic risk factors, and haplotypes of the monoamine oxidase (MAO) genes are among genetic variations increasing the risk of drug-related problems. However, data on the interactions between inheritable risk factors and early life stress (ES) with respect to predisposition to cocaine abuse are limited. Here, we show that a mouse model containing both genetic (low-activity alleles of the *MAO A* gene; MAOA^*Neo*^) and environmental (i.e., ES) variables displays a long lasting increased sensitivity to repeated *in vivo* cocaine psychomotor stimulant actions associated with a reduction of GABAA receptor-mediated inhibition of dopamine neurons of the ventral tegmental area (VTA). Depolarization-induced suppression of inhibition (DSI), a 2-arachidonoylglycerol (2-AG)-dependent form of short-term plasticity, also becomes readily expressed by dopamine neurons from MAOA^*Neo*^ES mice treated repeatedly with cocaine. Activation of either dopamine D2 or CB1 receptors is required for cocaine-induced DSI expression, decreased GABA synaptic efficacy, and hyperlocomotion. Next, *in vivo* pharmacological enhancement of 2-AG signaling during repeated cocaine exposure occludes its actions both *in vivo* and *ex vivo*. This data extends our knowledge of the multifaceted sequelae imposed by this gene by environment interaction in VTA dopamine neurons of male pre-adolescent mice, contributing to our understanding of neural mechanisms of vulnerability for early onset cocaine use disorder.

## Introduction

Substance use disorders are complex multifactorial conditions influenced by specific interactions among inheritable, biological, and environmental risk factors (Vink, 2016). Among the latter, exposure to early life stress (ES), such as child neglect, maltreatment, social isolation, and other traumas, represents an important environmental risk factor for the development of several neuropsychiatric disorders (e.g., depressive states, anxiety, substance use disorders) (Enoch, 2011; Hanson et al., 2021). Particularly, experiences of child abuse and maltreatment increase the likelihood of developing substance use disorders and lower the age of initial drug use (Lo Iacono et al., 2018; Scheidell et al., 2018; Santo et al., 2021). Early life adversity also influences psychostimulant-induced behavioral sensitization, a key component of drug addiction process (Bradberry, 2007; Koob and Volkow, 2010). Hence, animal studies show that ES, such as maternal separation, increases behavioral sensitization after repeated cocaine exposure (Delavari et al., 2016; Viola et al., 2016; Vannan et al., 2018).

An individual genetic background also contributes to the onset and the progression of the development of substance use disorders (Kreek et al., 2005; Fernandez-Castillo et al., 2022). Specific sets of genes involved in the mechanism of action of psychostimulants, particularly in the heavily-abused cocaine, result implicated in the development of this disorder, such as those related to monoamine (serotonin, dopamine, and norepinephrine) systems (Howell and Kimmel, 2008; Moeller et al., 2014). Among genetic factors, monoamine oxidase A (MAOA) gene plays an important role (Vanyukov et al., 2007; Stogner and CL, 2013). MAOA enzyme is a key enzyme involved in monoamine degradation. Low activity of MAOA gene has long been associated with drug dependence, particularly when one is exposed to early life adversity (Vanyukov et al., 2007; Stogner and CL, 2013; Fite et al., 2019a; Fite et al., 2019b). In humans, cocaine users with low activity of MAOA enzyme show an increased sensitivity to cocaine dependence across a lifetime, and this is associated with alterations in brain regions involved in reward processing, executive function and learning, which play a role in drug addiction (Vanyukov et al., 2004; Vanyukov et al., 2007; Alia-Klein et al., 2011; Verdejo-Garcia et al., 2013). The MAOA enzyme also plays an age-specific role in the etiopathogenesis of aggressive behavior (Yu et al., 2014a; Yu et al., 2014b), a trait that increases susceptibility for early onset of substance use (Brady et al., 1998; Miczek et al., 2004). Of note, a hypomorphic MAOA mouse (MAO^*Neo*^) when subjected to ES displays an aggressive behavior at pre-adolescence tied to a dysfunctional dopamine signaling (Frau et al., 2019a). Similarly, aberrant function of mesocorticolimbic dopamine pathway has been associated with cross-sensitization between early life stress and cocaine exposure (Brake et al., 2004; Enoch, 2011; Castro-Zavala et al., 2021) and to in

Despite this converging evidence, an unanswered question in cocaine research is whether the individuals with a low MAOA gene activity who experienced early life adversity display a biological predisposition toward the development of cocaine use. We, therefore, took advantage of a mouse model that of this gene by environment (G x E) interaction that is MAO^*Neo*^ ES mouse (Godar et al., 2019) and subjected it to discrete regimens of in vivo cocaine exposure. We found that MAO^*Neo*^ ES mice are vulnerable to psychostimulant effects of repeated in vivo cocaine exposure. This hyper-responsiveness is associated to persistent synaptic changes at inhibitory afferents onto dopamine cells in the ventral tegmental area (VTA). Such synaptic modifications require the recruitment of an endocannabinoid retrograde signal and the activation of dopamine D2 (DAD2) and type-1 cannabinoid (CB1) receptors, molecular targets exploited to prevent cocaine-induced sensitization.

## Results

### Repeated in vivo cocaine exposure cross-sensitizes with early life stress in hypomorphic MAOA mice

Low activity of MAOA uVNTR alleles in males predisposes to early onset of substance abuse (Vanyukov et al., 2004; Vanyukov et al., 2007). Hence, in the present study, we evaluated whether preadolescent (PND 28-30) MAOA^*Neo*^ mice subjected to ES could be more susceptible to psychostimulant effects of cocaine. As expected, in male pre-adolescent MAOA^*Neo*^ mice, a single in vivo exposure to cocaine (15 mg/kg, i.p.) increases locomotor activity, in terms of both horizontal counts and duration, irrespective of ES (Figure 1A,B). Next, repeated in vivo exposure to cocaine (7 days, once daily) unveils this G x E interaction, being MAOA^*Neo*^ES mouse hyperlocomotion enhanced when compared to the other experimental groups (Figure 1A,C). Notably, this hyper-responsiveness to cocaine persisted after one week of withdrawal (Figure 1A,D).

**Figure 1.**
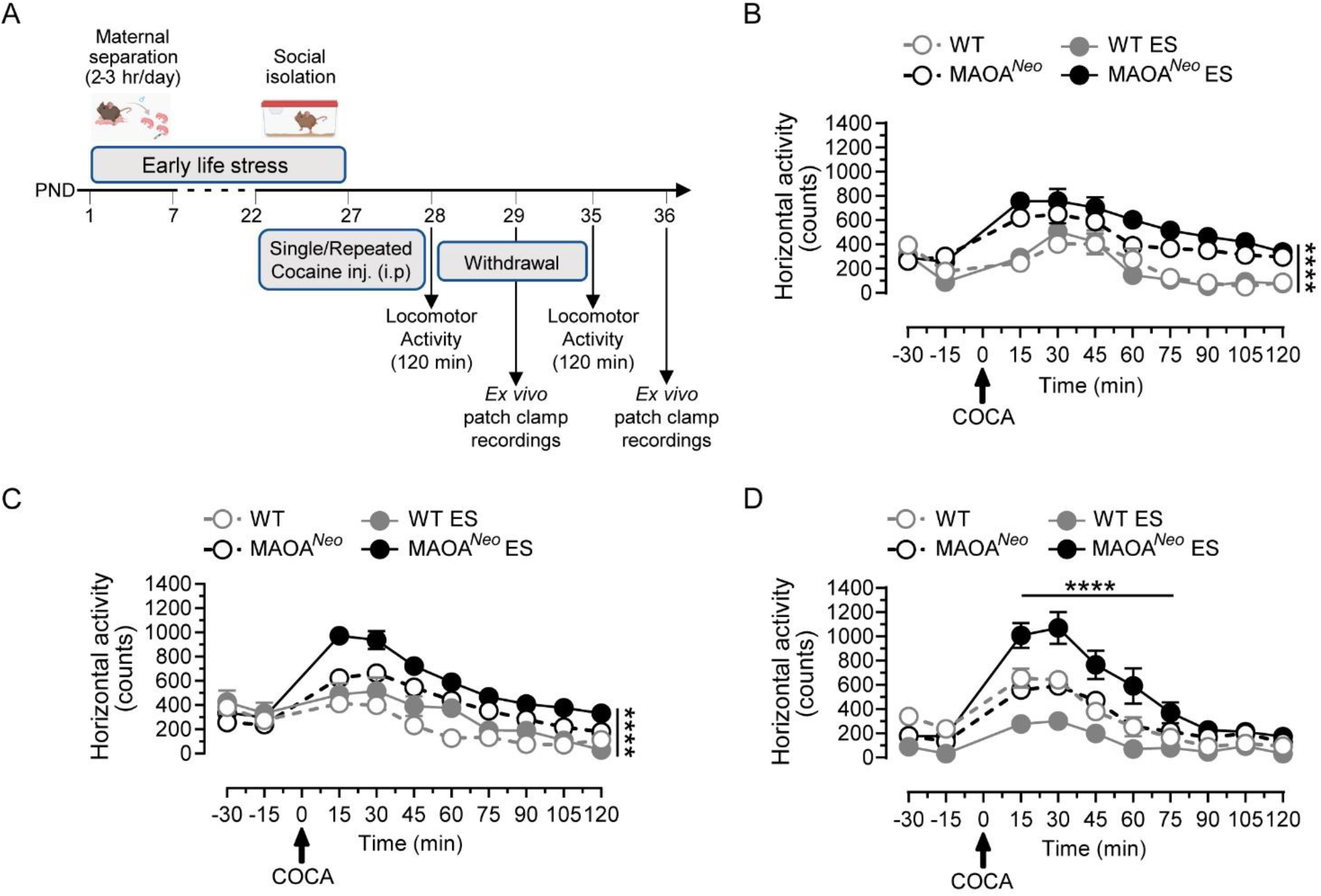
Psychostimulant effects of cocaine in MAOA^*Neo*^ mice subjected to early life stress. (**A**) Schematic experimental design to impose early life stress (ES) and to assess the psychostimulant response to cocaine after single and repeated exposure, as well as after 1 week of withdrawal. 24 hours after the last cocaine injection of repeated treatment and after 1 week of withdrawal, mice were subjected to whole-cell patch-clamp recordings. (**B**) Acute cocaine administration (COCA, 15 mg/kg, i.p.) increases the locomotor activity of MAOA^*Neo*^ mice, in terms of horizontal activity and duration (Two-way RM ANOVA followed by Tukey’s multiple comparisons test, Time effect F_(9,360)_=38,39, p<0,0001, Gene x ES effect F_(3,40)_=31,98, p<0,0001, Subject effect F_(40,360)_=3,46, p<0,0001, Interaction F_(27, 360)_=3,84, p<0,0001, n_mice_=10 WT and MAOA^*Neo*^, 11 WT ES, 13 MAOA^*Neo*^ ES). (**C**) Repeated exposure to cocaine for 1 week enhances the psychostimulant response to cocaine in MAOA^*Neo*^ ES mice (Two-way RM ANOVA followed by Tukey’s multiple comparisons test, Time effect F_(9,486)_=66,05, p<0,0001, Gene x ES effect F_(3,54)_=29,78, p<0,0001, Subject effect F_(54,486)_=5,12, p<0,0001, Interaction F_(27,486)_=7,65, p<0,0001, n_mice_=16 WT and MAOA^*Neo*^, 7 WT ES, 19 MAOA^*Neo*^ ES). (**D**) The enhanced locomotor activity persists long after 1 week of withdrawal in MAOA^*Neo*^ ES mice (Two-way RM ANOVA followed by Tukey’s multiple comparisons test, Time effect F_(10,320)_=54,76, p<0,0001, Gene x ES effect F_(3,32)_=12,01, p<0,0001, Subject effect F_(32,320)_=4,61, p<0,0001, Interaction F_(30,320)_=5,61, p<0,0001, n_mice_=10 WT, MAOA^*Neo*^, MAOA^*Neo*^ ES, 6 WT ES). Graphs show spontaneous locomotor (horizontal) activity measured as counts (bin=15 min) in a novel open field arena. Data are represented as mean ± SEM.

### Repeated in vivo cocaine exposure decreases GABAergic transmission onto dopamine cells

Repeated in vivo exposure to cocaine alters synaptic plasticity in the VTA, which plays an important role in behavioral consequences of cocaine use and abuse (Borgland et al., 2004; Bellone et al., 2021). To measure synaptic properties of VTA dopamine cells, we next performed whole-cell patch-clamp recordings in acute VTA slices 24 hr after the last cocaine administration. We first recorded AMPA excitatory postsynaptic currents (EPSCs) from putative dopamine neurons and found no differences among groups in current-voltage relationships (I-V) (Figure 2A) and in the AMPA/NMDA ratio (Figure 2B). Statistical analysis revealed a main effect of ES in the paired-pulse modulation (EPSC2/EPSC1) of AMPA EPSCs (Figure 2C) and in the decay time of NMDA EPSCs (Figure 2D). Altogether, these data suggest that changes at excitatory inputs onto dopamine cells do not account for the enhanced responsiveness to cocaine exhibited exclusively by MAOA^*Neo*^ ES mice upon repeated in vivo exposure to cocaine.

**Figure 2.**
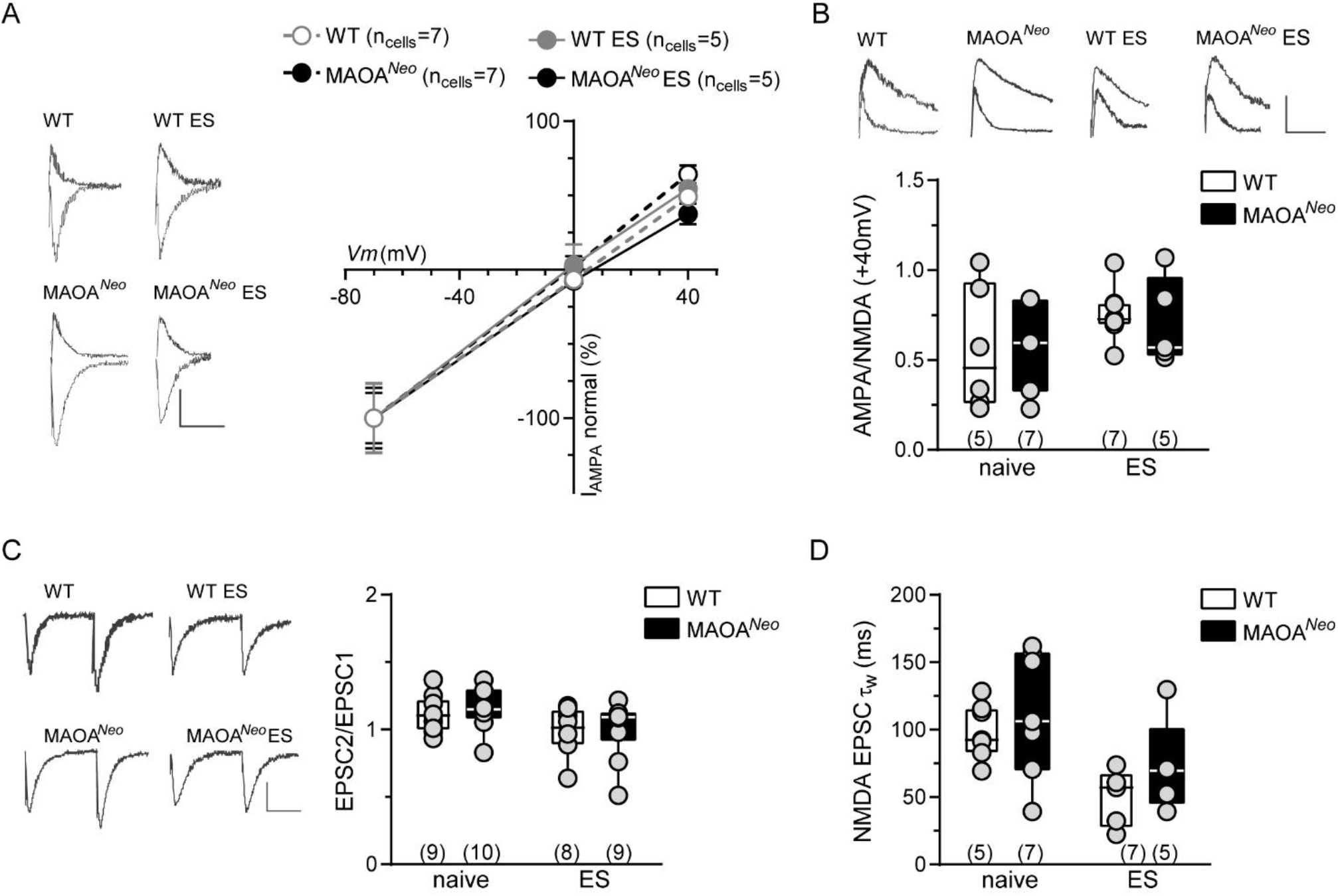
Effect of repeated cocaine exposure to excitatory synaptic properties of VTA DA neurons in MAOA^*Neo*^ mice subjected to ES. (**A**) Current-voltage relationship (I–V) curves of AMPA EPSCs recorded from DA neurons 24 hr after the last injection of repeated cocaine exposure (1 week) in WT (n_cells,mice_ =7,5), MAOA^*Neo*^ (n_cells,mice_=7,7), WT ES (n_cells,mice_=6,7) and MAOA^*Neo*^ ES (n_cells,mice_=5,5) mice (Two-way ANOVA, Gene x ES effect F_(3,63)_=0,42, p=0,739, mV effect F_(2,63)_=118,1, p<0,0001, Interaction F_(6,63)_=0,17, p=0,98). Insets show representative traces of AMPA EPSCs recorded at −70 mV and +40 mV. Calibration bar: 25 ms, 50 pA. Each symbol represents the averaged value (± SEM) obtained from different cells. (**B**) The AMPA/NMDA ratio is not affected by repeated cocaine exposure in WT (n_cells,mice_=6,5), MAOA^*Neo*^ and WT ES (n_cells,mice_=7,7 per group), MAOA^*Neo*^ ES (n_cells,mice_=5,5) mice (Two-way ANOVA, Gene effect F_(1,21)_=0,04, p=0,85, ES effect F_(1,21)_=2,86, p=0,105, Interaction F_(1,21)_=0,11, p=0,743). Insets show representative traces of AMPA, and NMDA EPSCs traces recorded from DA neurons held at +40mV in VTA slices. Calibration bar: 25 ms, 50 pA. (**C**) Graph panel summarizing the effect of ES on paired-pulse ratio (EPSC2/EPSC1) of AMPA EPSCs recorded from WT (n_cells,mice_=9,9), MAOA^*Neo*^ (n_cells,mice_=10,10), WT ES (n_cells,mice_=8,8) and MAOA^*Neo*^ ES (n_cells,mice_=10,9) slices 24 hr after the end of repeated cocaine exposure (Two-way ANOVA, Gene effect F_(1,33)_=0,28, p=0,597, ES effect F_(1,33)_=5,66, p=0,023, Interaction F_(1,33)_=0,02, p=0,89). Insets show representative traces of paired AMPA EPSCs. Calibration bar: 25 ms, 50 pA. (**D**) Quantification of the data showing NMDAR EPSC decay time kinetics (weighted tau, τ) in WT (n_cells,mice_=7,5), MAOA^*Neo*^ (n_cells,mice_=7,7), WT ES (n_cells,mice_=5,7) and MAOA^*Neo*^ ES (n_cells,mice_=5,5) slices (Two-way ANOVA, Gene effect F_(1,20)_=1,69, p=0,208, ES effect F_(1,20)_=10,39, p=0,004, Interaction F_(1,20)_=0,134, p=0,718). Unless otherwise indicated, graphs show box- and-whisker plots (including minima, maxima and median values, and lower and upper quartiles) with each circle representing a single cell recorded (numbers in the brackets represent number of the mice).

Next, we examined the properties of GABA synaptic inputs on VTA dopamine neurons by recording GABAA inhibitory postsynaptic currents (IPSCs), since repeated in vivo cocaine exposure also diminishes GABA transmission in rat midbrain dopamine neurons (Liu et al., 2005; Pan et al., 2008b). MAOA^*Neo*^ ES mouse dopamine cells exhibited a decreased maximal amplitude of GABAA IPSCs (Figure 3A), an effect accompanied by a facilitation of GABAA IPSCs (IPSC2/IPSC1) (Figure 3B) and that lasted after one week from the last administration (Figure 3C, D). Since basal inhibitory synaptic properties of VTA dopamine neurons of MAOA^*Neo*^ mice are not affected by ES (data not shown), this reduced GABA efficacy on dopamine cells could be ascribed to a higher responsiveness to cocaine displayed by MAOA^*Neo*^ ES dopamine cells. Accordingly, cocaine is more potent and more effective in decreasing GABAA IPSC amplitude in MAOA^*Neo*^ ES mouse dopamine neurons (Figure 3E). In rats, cocaine-induced reduction of inhibitory strength on dopamine neurons depends on the recruitment of endocannabinoid system mediating a long-but not a short-term form of synaptic plasticity (Pan et al., 2008b). Nonetheless, a short-term form of synaptic plasticity termed depolarization-induced suppression of inhibition (DSI) is associated to vulnerability to drugs of abuse (Melis et al., 2013; Melis et al., 2014). Therefore, we examined whether cocaine-exposed MAOA^*Neo*^ ES mouse dopamine cells could express DSI at these synapses by phasically activating endocannabinoid signaling via somatic current injection into dopamine cells (from -70 to +40 mV, 3 s). MAOA^*Neo*^ ES mouse dopamine neurons did not express DSI (data not shown) unless subjected to repeated in vivo cocaine administration (Figure 4A, B). DSI was accompanied by an increased facilitation of these synapses (Figure 4C), suggestive of a decreased probability of GABA release (Melis et al., 2002). Since DSI expression persisted after one week of withdrawal (Figure 4D-F), this plasticity might contribute to the hyperresponsiveness displayed by MAOA^*Neo*^ ES mice to repeated in vivo cocaine exposure.

**Figure 3.**
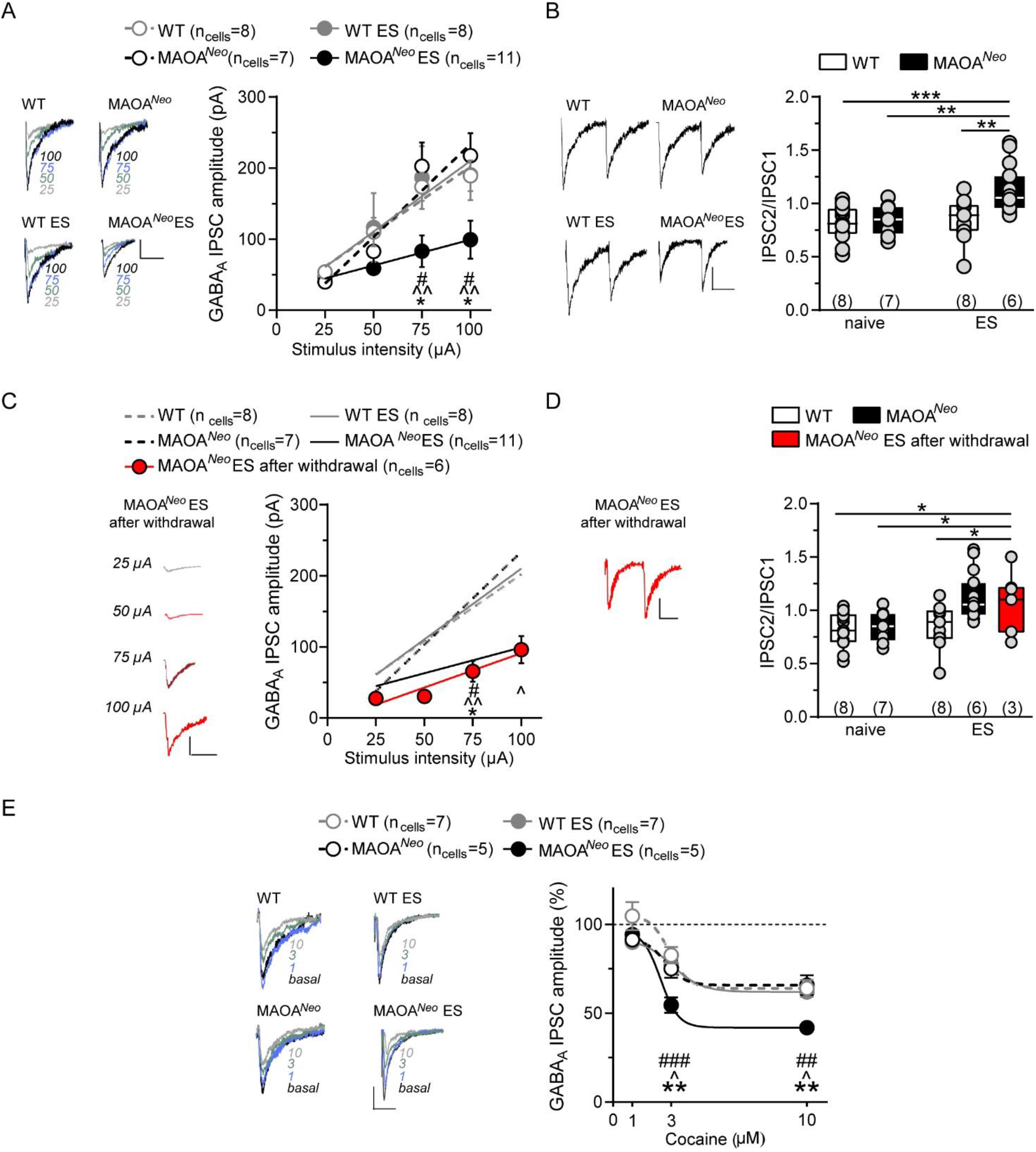
Effect of repeated cocaine exposure to inhibitory transmission on VTA DA neurons in MAOA^*Neo*^ mice subjected to ES. (**A**) Input–output relationships of GABAA-mediated IPSCs in WT and WT ES (n_cells,mice_=8,8 per group), MAOA^*Neo*^ (n_cells,mice_ =7,7) and MAOA^*Neo*^ ES (n_cells,mice_=11,6) mice recorded 24 hr after the last treatment of repeated cocaine exposure (Two-way RM ANOVA, Stimulus intensity effect F_(3,90)_=43,91, p<0,0001, Gene x ES effect F_(3,30)_=2,87, p=0,053, Subject effect F_(30,90)_=4,97, p<0,0001, Interaction F_(9,90)_=2,69, p=0,008, Tukey’s multiple comparisons post hoc test, 75 μA, 100 μA, ^#^MAOA^*Neo*^ ES vs WT, p<0,05, ^^MAOA^*Neo*^ ES vs MAOA^*Neo*^, p<0,01, *MAOA^*Neo*^ ES vs WT ES, p<0,05). Each symbol represents the averaged value (± SEM) obtained from different cells. Insets show representative traces of GABAA IPSC recorded at each stimulus intensity. Calibration bar: 25 ms, 50 pA. (**B**) Graph panel summarizing the averaged paired-pulse ratio (IPSC2/IPSC1) of GABAA IPSCs for all cells recorded 24 hr after the last treatment of repeated cocaine exposure in WT and WT ES (n_cells,mice_=8,8 per group), MAOA^*Neo*^ (n_cells,mice_ =7,7) and MAOA^*Neo*^ ES (n_cells,mice_=10,6) mice (Two-way ANOVA, Gene effect F_(1,47)_=7,53, p=0,0086, ES effect F_(1,47)_=9,05, p=0,004, Interaction F_(1,47)_=5,46, p=0,024, Tukey’s multiple comparisons post hoc test, ***MAOA^*Neo*^ ES vs WT, p=0,0003, **MAOA^*Neo*^ ES vs MAOA^*Neo*^, p=0,003, **MAOA^*Neo*^ ES vs WT ES, p=0,0017). Inset shows representative traces of paired GABAA IPSCs recorded from VTA putative DA neurons. Calibration bar: 25 ms, 50 pA. (**C**) Input–output relationships of GABAA-mediated IPSCs shows that repeated cocaine effect in reducing synaptic efficacy of these synapses persists after 1 week of withdrawal in MAOA^*Neo*^ ES mice (n_cells,mice_=6,3) (Two-way RM ANOVA, Stimulus intensity effect F_(3,105)_=46,72, p<0,0001, Gene x ES effect F_(4,35)_=3,97, p=0,009, Subject effect F_(35,105)_=4,92, p<0,0001, Interaction F_(12,105)_=2,71, p=0,003, Tukey’s multiple comparisons post hoc test, 75 μA, ^#^MAOA^*Neo*^ ES after withdrawal vs WT, p<0,05, ^^MAOA^*Neo*^ ES after withdrawal vs MAOA^*Neo*^, p<0,01, *MAOA^*Neo*^ ES after withdrawal vs WT ES, p<0,05, 100 μA, ^MAOA^*Neo*^ ES after withdrawal vs MAOA^*Neo*^, p<0,05). Each symbol represents the averaged value (± SEM) obtained from different cells. Insets show representative traces of IPSC recorded at each stimulus intensity for MAOA^*Neo*^ ES after 1 week of withdrawal. Calibration bar: 25 ms, 50 pA. (**D**) Graph panel displays that the facilitation in the averaged paired-pulse ratio (IPSC2/IPSC1) of GABAA IPSCs lasts for 1 week of withdrawal in MAOA^*Neo*^ ES (n_cells,mice_=6,3) mice (Two-way ANOVA, Gene effect F_(2,56)_=7,11, p=0,0018, ES effect F_(1,56)_=8,58, p=0,004, Interaction F_(2,56)_=2,73, p=0,073, Tukey’s multiple comparisons post hoc test, *MAOA^*Neo*^ ES after withdrawal vs WT, p=0,013, *MAOA^*Neo*^ ES after withdrawal vs MAOA^*Neo*^, p=0,04, *MAOA^*Neo*^ ES after withdrawal vs WT ES, p=0,03). Inset shows representative trace of paired GABAA IPSCs recorded from VTA putative DA neurons for MAOA^*Neo*^ ES after 1 week of withdrawal. Calibration bar: 25 ms, 50 pA. (**E**) Dose–response curves for the inhibition of the GABAA IPSCs by bath-application of cocaine (1, 3, 10 μM) displays a larger effect in MAOA^*Neo*^ ES (n_cells,mice_=5,5) than in WT (n_cells,mice_=7,7), MAOA^*Neo*^ (n_cells,mice_=5,5) and WT ES (n_cells,mice_=7,6) cells recorded 24 hr after the last injection of repeated cocaine exposure (Two-way RM ANOVA, Cocaine concentration effect F_(2,40)_=71,07, p<0,0001, Gene x ES effect F_(3,20)_=6,965, p=0,002, Subject effect F_(20,40)_=1,54, p=0,12, Interaction F_(6,40)_=2,53, p=0,0358, Tukey’s multiple comparisons post hoc test, 3μM, ^###^MAOA^*Neo*^ ES vs WT, p<0,001, ^MAOA^*Neo*^ ES vs MAOA^*Neo*^, p<0,05, **MAOA^*Neo*^ ES vs WT ES, p<0,01, 10μM, ^##^MAOA^*Neo*^ ES vs WT, p<0,01, ^^MAOA^*Neo*^ ES vs MAOA^*Neo*^, p<0,01, *MAOA^*Neo*^ ES vs WT ES, p<0,05. Insets show IPSC traces recorded after 1 μM (blue), 3 μM (green), and 10 μM (grey) cocaine application. Calibration bar, 50 ms, 100 pA. Each symbol represents the averaged value (± SEM) obtained from different cells. Unless otherwise indicated, graphs show box- and-whisker plots (including minima, maxima and median values, and lower and upper quartiles) with each circle representing a single cell recorded (numbers in the brackets represent number of the mice).

**Figure 4.**
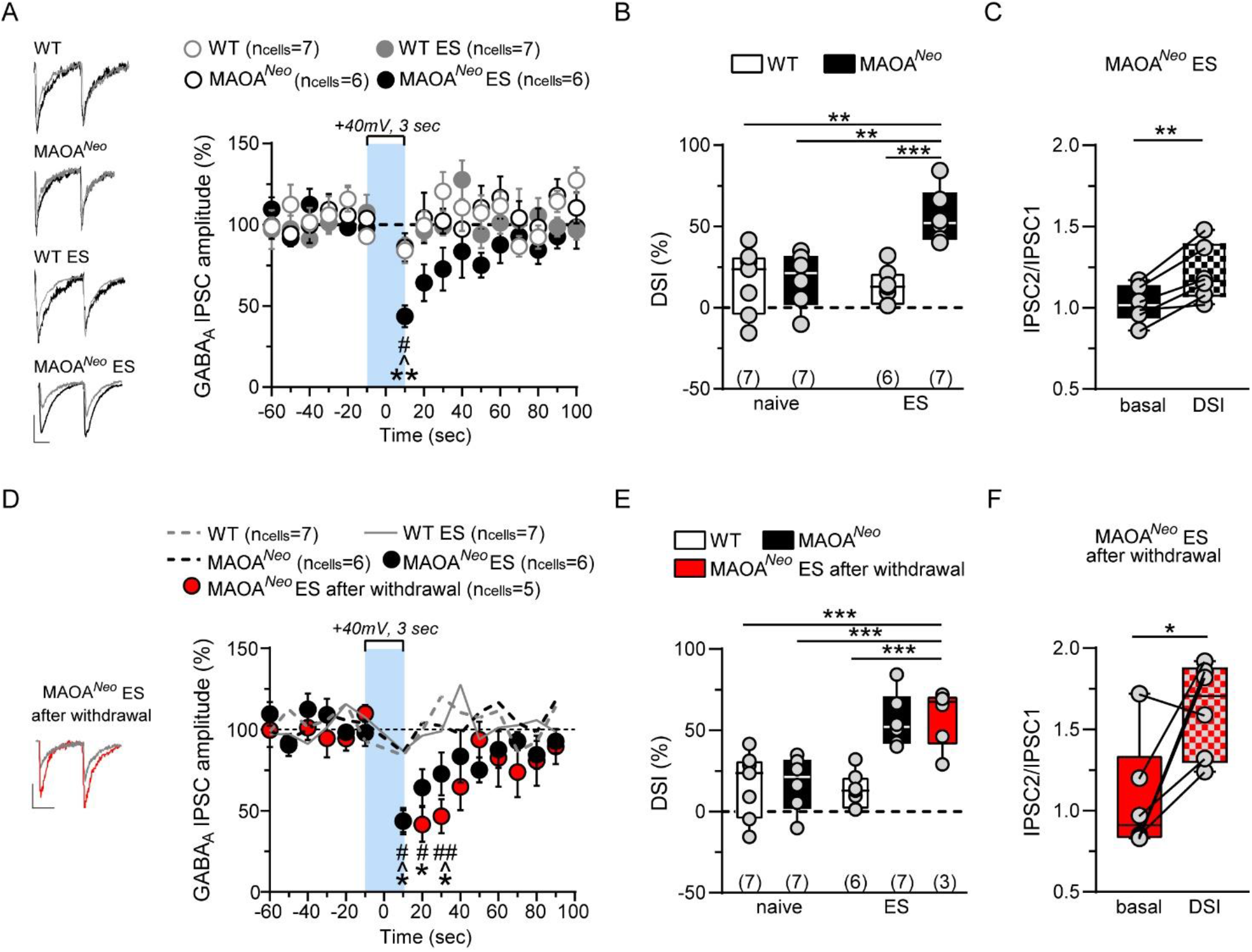
Endocannabinoid modulation of inhibitory transmission arising from rostral afferents is shown by MAOA^*Neo*^ mice subjected to ES after repeated cocaine exposure. (**A**) Graph panel showing the time course of DSI. Application of a depolarizing pulse from -70 to +40 mV for 3 sec on VTA DA neurons induces reduction of GABAA IPSCs evoked by stimulating rostral afferents in MAOA^*Neo*^ ES mice recorded 24 hr after the end of repeated cocaine treatment (Two way RM ANOVA, Time effect F_(6.96,153)_=3,787, p=0,0008, Gene x ES effect F_(3,22)_=3,895, p=0,023, Subject effect F_(22,440)_=2,61, p=0,0001, Interaction F_(60,440)_=1,092, p=0,306, Tukey’s multiple comparisons post hoc test, 10 s, ^#^MAOA^*Neo*^ ES vs WT, p=0,011, ^MAOA^*Neo*^ ES vs MAOA^*Neo*^, p=0,016, **MAOA^*Neo*^ ES vs WT ES, p=0,002). GABAA IPSCs amplitude was normalized to the averaged value (dotted line) before depolarization. Each symbol represents the averaged value (± SEM) obtained from different cells, WT (n_cells,mice_=7,7), WT ES (n_cells,mice_=7,6), MAOA^*Neo*^ and MAOA^*Neo*^ ES (n_cells,mice_=6,7 per group). Left panel shows representative traces of IPSC recorded before (black) and after DSI (grey). Calibration bar, 25 ms, 50 pA. (**B**) Averaged data for DSI induced by depolarizing pulse with a duration of 3 sec are plotted (Two way ANOVA, ES effect F_(1,22)_=7,4, p=0,013, Gene effect F_(1,22)_=11,21, p=0,003, Interaction F_(1,22)_=10,25, p=0,004, Tukey’s multiple comparisons post hoc test, **MAOA^*Neo*^ ES vs WT, p=0,002, **MAOA^*Neo*^ ES vs MAOA^*Neo*^, p=0,003, ***MAOA^*Neo*^ ES vs WT ES, p<0,001). (**C**) Bar graph summarizing the averaged paired-pulse ratio (IPSC2/IPSC1) of rostral GABAA IPSCs for all cells recorded in MAOA^*Neo*^ ES mice before (basal) and after (DSI) depolarization pulse (Two tailed paired t-test, t_(5)_=4,858, **p=0,0046). (**D**) Graph panel showing the time course of DSI after 1 week of withdrawal. Application of a depolarizing pulse from -70 to +40 mV for 3 sec on VTA DA neurons induces reduction of GABAA IPSCs evoked by stimulating rostral afferents in MAOA^*Neo*^ ES after withdrawal (n_cells,mice_=5,3) mice (Two way RM ANOVA, Time effect F_(7.559,196.5)_=5,20, p<0,0001, Gene x ES effect F_(4,26)_=8,07, p=0,0002, Subject effect F_(26,364)_=2,07, p=0,0019, Interaction F_(56,364)_=1,64, p=0,0042, Tukey’s multiple comparisons post hoc test, 10 s, ^#^MAOA^*Neo*^ ES after withdrawal vs WT, p=0,035, ^MAOA^*Neo*^ ES after withdrawal vs MAOA^*Neo*^, p=0,038, *MAOA^*Neo*^ ES after withdrawal vs WT ES, p=0,018, 20 s, ^#^MAOA^*Neo*^ ES after withdrawal vs WT, p=0,017, *MAOA^*Neo*^ ES after withdrawal vs WT ES, p=0,019, 30 s, ^##^MAOA^*Neo*^ ES after withdrawal vs WT, p=0,006, ^MAOA^*Neo*^ ES after withdrawal vs MAOA^*Neo*^, p=0,018, *MAOA^*Neo*^ ES after withdrawal vs WT ES, p=0,024). GABAA IPSCs amplitude was normalized to the averaged value (dotted line) before depolarization. Each symbol represents the averaged value (± SEM) obtained from different cells. Left panel shows representative traces of IPSC recorded before (red) and after DSI (grey). Calibration bar, 25 ms, 50 pA. (**E**) Averaged data for DSI induced by depolarizing pulse with a duration of 3 sec are plotted for all cells recorded in MAOA^*Neo*^ ES mice after 1 week of withdrawal (Two way ANOVA, ES effect F_(1,32)_=39,19, p<0,0001, Gene x treatment effect F_(2,32)_=6,90, p=0,003, Interaction F_(2,32)_=13,70, p<0,0001, Tukey’s multiple comparisons post hoc test, ***MAOA^*Neo*^ ES after withdrawal vs WT, p=0,0003, ***MAOA^*Neo*^ ES after withdrawal vs MAOA^*Neo*^, p=0,0008, ***MAOA^*Neo*^ ES vs WT ES, p=0,0001). (**F**) Bar graph summarizing the averaged paired-pulse ratio (IPSC2/IPSC1) of rostral GABAA IPSCs for all cells recorded in MAOA^*Neo*^ ES mice after 1 week of withdrawal before (basal) and after (DSI) depolarization pulse (Two tailed paired t-test, t_(5)_=3,094, *p=0,027). Each symbol represents the averaged value (± SEM) obtained from different cells. Unless otherwise indicated, graphs show box- and-whisker plots (including minima, maxima and median values, and lower and upper quartiles) with each circle representing a single cell recorded (numbers in the brackets represent number of the mice).

Metaplastic changes in the molecular architecture of endocannabinoid system occurring in MAOA^*Neo*^ ES mice repeatedly exposed in vivo to cocaine could explain the expression of DSI. To test whether CB1 receptor function and/or number would be affected by repeated in vivo exposure to cocaine, we built a concentration-response relationship for the CB1R/CB2R agonist WIN55,212-2 (WIN; 0.01-3 μM), but found no differences among the experimental groups (Figure 5A). The half-life of the endocannabinoid 2-arachidonoylglycerol (2-AG) is often the rate-limiting step that determines the time course of DSI in the brain (Pan et al., 2009), including at inhibitory synapses in the VTA (Melis et al., 2013; Melis et al., 2014). 2-AG half-life is tightly regulated by its main degrading enzyme monoacylglycerol lipase (MAGL) (Blankman et al., 2007). We, therefore, examined the effects of the MAGL inhibitor JZL184 on DSI. In the presence of JZL184 (100 nM, 5 min pre-incubation), which per se was ineffective (Figure 5B), all the experimental groups expressed similar DSI and paired pulse facilitation (Figure 5C,D).

**Figure 5.**
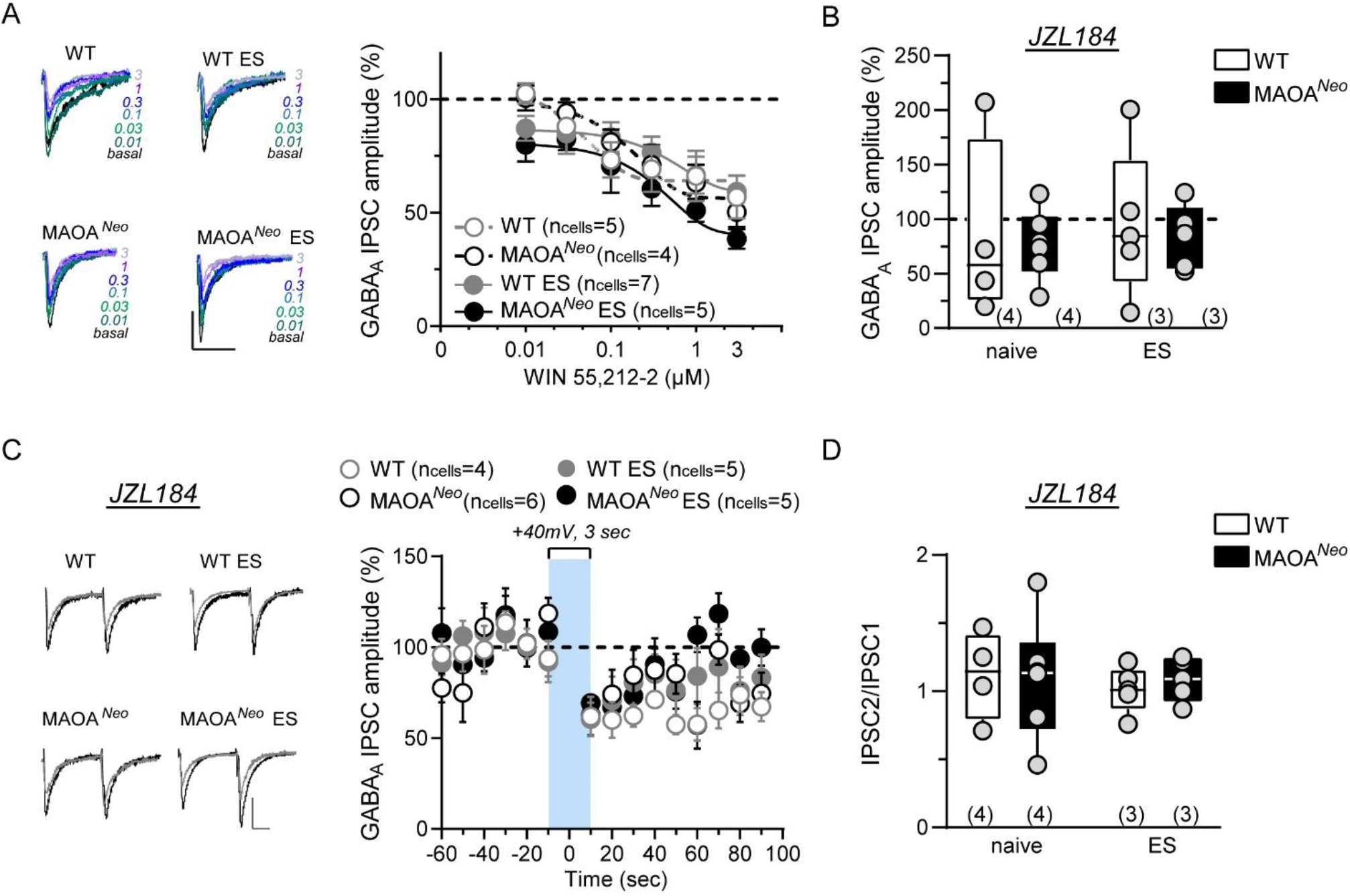
CB1 receptors activation and MAGL inactivation produce similar effects among groups on VTA DA neurons after repeated cocaine exposure. (**A**) Dose–response curves for percentage inhibition in GABAA IPSCs amplitude by the CB1R agonist WIN55,212–2 as recorded from VTA DA cells and evoked by stimulating rostral afferents in WT (n_cells,mice_=5,4), MAOA^*Neo*^ (n_cells,mice_=4,2), WT ES (n_cells,mice_=7,6) and MAOA^*Neo*^ ES (n_cells,mice_=5,5) (Two way RM ANOVA, Gene x ES effect F_(3,18)_=1,27, p=0,364, Concentration effect F_(3.02,52.6)_=34,58, p<0,0001, Interaction F_(15,87)_=1,0, p=0,459. Each symbol represents the averaged value (± SEM) obtained from different cells. The insets show IPSCs before (black), and during perfusion of WIN (0,01-3 μM). Calibration bar, 25 ms, 100 pA. (**B**) Bar graph displays the effect of MAGL inactivation in the amplitude of GABAA IPSCs for all cells recorded in WT (n_cells,mice_=4,4), MAOA^*Neo*^ (n_cells,mice_=6,4), WT ES and MAOA^*Neo*^ ES (n_cells,mice_=5,3 per group) mice (Two way ANOVA, ES effect F_(1,16)_=0,109, p=0,745, Gene effect F_(1,16)_=0,183, p=0,674, Interaction F_(1,16)_=0,003, p=0,953). (**C**) Graph panel showing the time course of DSI in the presence of JZL184 (100 nM) in WT (n_cells,mice_=4,4), MAOA^*Neo*^ (n_cells,mice_=5,4), WT ES (n_cells,mice_=6,3) and MAOA^*Neo*^ ES (n_cells,mice_=5,3). JZL184 induced similar DSI among groups. GABAA IPSCs amplitude was normalized to the averaged value (dotted line) before depolarization (Two way RM ANOVA, Time effect F_(6.64,106.2)_=6,78, p<0,0001, Gene x ES effect F_(3,16)_=2,43, p=0,102, Subject effect F_(16,224)_=1,85, p=0,025, Interaction F_(42,224)_=0,98, p=0,496). Each symbol represents the averaged value (± SEM) obtained from different cells. The insets show representative traces of IPSC recorded before (black) and after DSI (grey). Calibration bar, 25 ms, 50 pA (**D**) Bar graph displays that MAGL inactivation produced no differences in paired-pulse ratio (IPSC2/IPSC1) of rostral GABAA IPSCs for all cells recorded in WT (n_cells,mice_=4,4), MAOA^*Neo*^ (n_cells,mice_=6,4), WT ES and MAOA^*Neo*^ ES (n_cells,mice_=5,3 per group) mice (Two way ANOVA, ES effect F_(1,16)_=0,158, p=0,69, Gene effect F_(1,16)_=0,029, p=0,865, Interaction F_(1,16)_=0,13, p=0,725). Unless otherwise indicated, graphs show box- and-whisker plots (including minima, maxima and median values, and lower and upper quartiles) with each circle representing a single cell recorded (numbers in the brackets represent number of the mice).

### Dopamine and 2-AG co-participate to depress GABAergic inputs on dopamine neurons

In the rat VTA, dopamine D2 (DAD2) receptor activation is implicated in 2-AG synthesis and release from dopamine cells (Melis et al., 2004) as well as in cocaine-induced inhibition of synaptic strength of GABA inputs (Pan et al., 2008b). We, therefore, investigated whether similar mechanisms were involved in cocaine-induced effects in MAOA^*Neo*^ ES mice. Pharmacological in vivo pretreatment with the DAD2 receptor antagonist sulpiride (10 mg/kg, i.p., 7 days, 20 min before cocaine administration) resumed GABAergic transmission on VTA dopamine cells (Figure 6A,B), abolished DSI and its related paired-pulse facilitation (Figure 6C-E) and prevented cocaine-induced hyperlocomotion in MAOA^*Neo*^ ES mice (Figure 6F).

**Figure 6.**
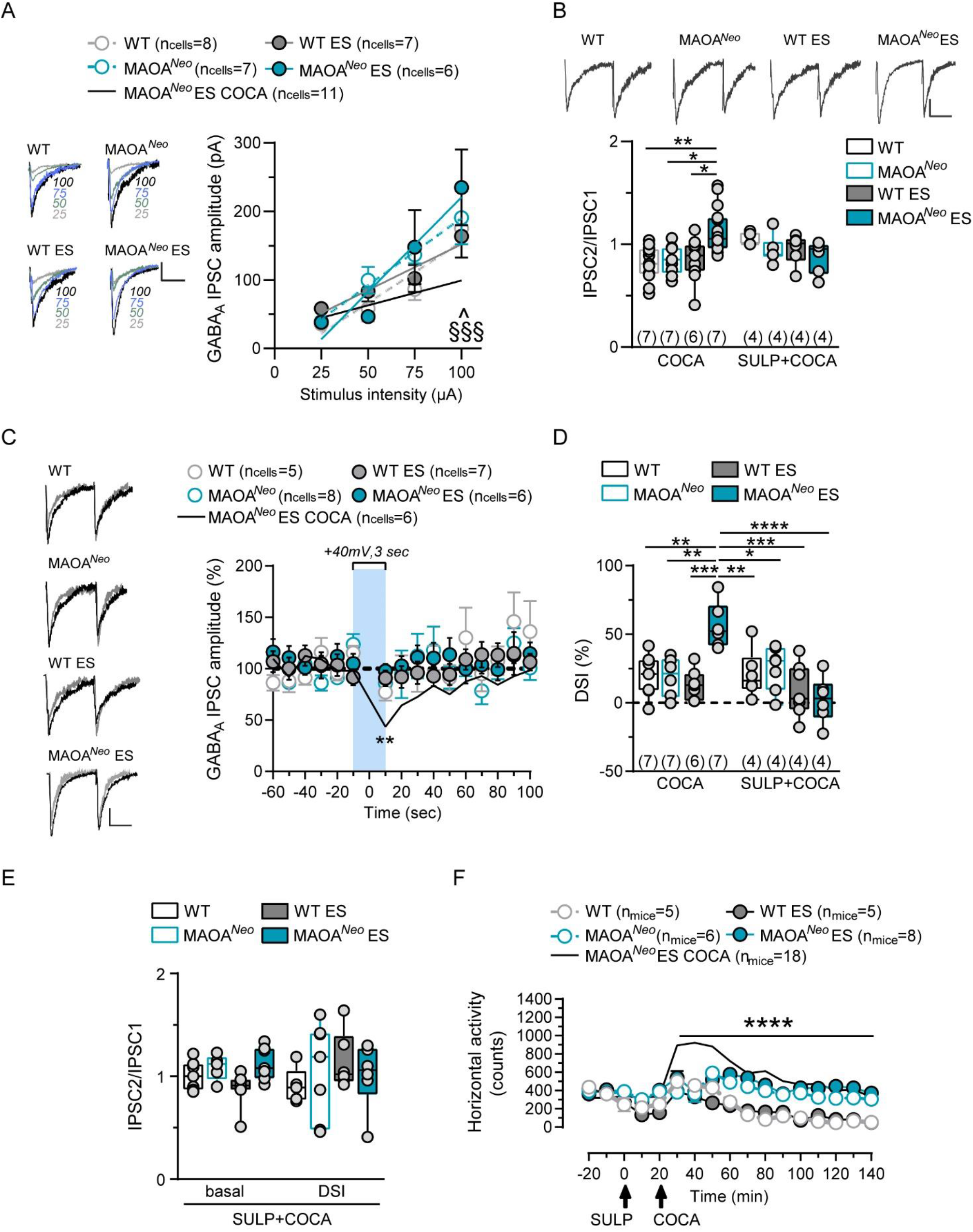
*In vivo* pretreatment with D2DR antagonist sulpiride restores synaptic properties and plasticity of inhibitory transmission on VTA DA neurons and prevents cocaine-induced hyperlocomotion in MAOA^*Neo*^ mice subjected to ES. (**A**) Input–output relationships of GABAA-mediated IPSCs shows the effect of pretreatment with DA2DR antagonist, sulpiride (SULP, 10 mg/kg, i.p) 20 m before each intraperitoneal cocaine injection (COCA, 15 mg/kg) on midbrain DA neurons 24 hrs after the last repeated treatment (1 week). SULP abolishes cocaine-induced reduction in the amplitude of GABAA IPSCs in MAOA^*Neo*^ ES mice and differences among groups, WT (n_cells,mice_=8,4), MAOA^*Neo*^ and WT ES (n_cells,mice_=7,4 per group) and MAOA^*Neo*^ ES (n_cells,mice_=6,4) mice (Two-way RM ANOVA, Stimulus intensity effect F_(3,99)_=45,22, p<0,0001, Gene x ES x Trt effect F_(4,33)_=1,53, p=0,215, Subject effect F_(33,99)_=3,19, p<0,0001, Interaction F_(12,99)_=2,38, p=0,0096, Tukey’s multiple comparisons post hoc test, 100 μA, ^MAOA^*Neo*^ ES COCA vs MAOA^*Neo*^ SULP+COCA, p<0,05, ^§§§^MAOA^*Neo*^ ES COCA vs MAOA^*Neo*^ ES SULP+COCA, p<0,001). Each symbol represents the averaged value (± SEM) obtained from different cells. Insets show representative traces of GABAA IPSCs recorded at each stimulus intensity. Calibration bar: 25 ms, 50 pA. (**B**) Graph panels summarizing the effect of pretreatment with SULP on averaged paired-pulse ratio (IPSC2/IPSC1) for DA cells recorded in WT (n_cells,mice_=8,4), MAOA^*Neo*^ and WT ES (n_cells,mice_=7,4 per group) and MAOA^*Neo*^ ES (n_cells,mice_=6,4) mice (Two way ANOVA, Gene x ES effect F_(3,66)_=1,33, p=0,271, Treatment effect F_(1,66)_=1,10, p=0,298, Interaction F_(3,66)_=5,72, p=0,0015, Tukey’s multiple comparisons post hoc test, ***MAOA^*Neo*^ ES COCA vs WT COCA, p=0,0003, **MAOA^*Neo*^ ES COCA vs MAOA^*Neo*^ COCA, p=0,0045, **MAOA^*Neo*^ ES COCA vs WT ES COCA, p=0,002). Inset shows representative traces of paired GABAA IPSCs recorded from VTA putative DA neurons. Calibration bar: 25 ms, 50 pA. (**C**) Graph panel shows the effect of pretreatment of sulpiride on time course of DSI 24 hr after the last repeated SULP+COCA treatment. Application of a depolarizing pulse from -70 to +40 mV for 3 sec on VTA DA neurons prevents DSI of GABAA IPSCs evoked by stimulating rostral afferents in WT (n_cells,mice_=5,4), MAOA^*Neo*^ (n_cells,mice_=8,4), WT ES (n_cells,mice_=7,4) and MAOA^*Neo*^ ES mice (n_cells,mice_=6,4) (Two way RM ANOVA, Time effect F_(6.77,183)_=3,22, p=0,003, Gene x ES effect F_(4,27)_=1,19, p=0,336, Subject effect F_(27,405)_=5,55, p<0,0001, Interaction F_(60,405)_=1,21, p=0,127, Tukey’s multiple comparisons post hoc test, 10 s, **p=0,0016 MAOA^*Neo*^ ES vs MAOA^*Neo*^ ES COCA). GABAA IPSCs amplitude was normalized to the averaged value (dotted line) before depolarization. Each symbol represents the averaged value (± SEM) obtained from different cells. Panel shows representative traces of IPSCs recorded before and after DSI (grey). Calibration bar, 25 ms, 50 pA. (**D**) Averaged data for DSI induced by depolarizing pulse with a duration 3 sec are plotted (Two way ANOVA, Gene x ES effect F_(3,44)_=2,80, p=0,050, Treatment effect F_(1,44)_=7,99, p=0,0071, Interaction F_(3,44)_=9,24, p<0,0001, Tukey’s multiple comparisons post hoc test, **MAOA^*Neo*^ ES COCA vs WT COCA, p=0,0038, **MAOA^*Neo*^ ES COCA vs MAOA^*Neo*^ COCA, p=0,0049, ***MAOA^*Neo*^ ES COCA vs WT ES COCA, p=0,0004, **MAOA^*Neo*^ ES COCA vs WT SULP+COCA, p=0,009, *MAOA^*Neo*^ ES COCA vs MAOA^*Neo*^ SULP+COCA, p=0,020, ***MAOA^*Neo*^ ES COCA vs WT ES SULP+COCA, p=0,0001, ****MAOA^*Neo*^ ES COCA vs MAOA^*Neo*^ ES SULP+COCA, p<0,0001). (**E**) Bar graph summarizing the averaged paired-pulse ratio (IPSC2/IPSC1) of GABAA IPSCs for all cells recorded after pretreatment with SULP, before (basal) and after (DSI) depolarization pulse (Two way ANOVA, Gene x ES effect F_(3,42)_=0,526, p=0,666, Basal vs DSI effect F_(1,42)_=0,004, p=0,949, Interaction F_(3,42)_=1,48, p=0,233). (**F**) Pretreatment of mice with DA2DR antagonist, sulpiride, for 1 week abolishes the cocaine-induced hyperlocomotion in MAOA^*Neo*^ mice exposed to ES (Two-way RM ANOVA followed by Tukey’s multiple comparisons test, Time effect F_(4.069, 158.7)_=11,48, p<0,0001, Gene x ES x treatment effect F_(4, 39)_ =11,73, p<0,0001, Subject effect F_(39, 624)_=7,87, p<0,0001, Interaction F_(64, 624)_=3,55, p<0,0001, n mice=5 WT, 6 MAOA^*Neo*^, 7 WT ES, 8 MAOA^*Neo*^ ES, 18 MAOA^*Neo*^ ES COCA). Graph shows spontaneous locomotor (horizontal) activity measured as counts (bin=10 min) before and after drugs injection at the last day of repeated treatment in a novel open field arena. Data are represented as mean ± SEM. Unless otherwise indicated, graphs show box- and-whisker plots (including minima, maxima and median values, and lower and upper quartiles) with each circle representing a single cell recorded (numbers in the brackets represent number of the mice).

Notably, DAD2 and CB1 receptors target the same downstream signaling pathway to produce long term depression at GABAergic synapses on dopamine neurons (Pan et al., 2008a). Next, we pharmacologically blocked in vivo CB1 receptors by administering the potent antagonist AM281 (2.5 mg/kg, i.p. 7 days) during cocaine exposure (20 min before each cocaine injection). In MAOA^*Neo*^ ES mice, in vivo AM281 pre-treatment averted the decreased GABA synaptic efficacy, the differences in paired-pulse modulation (Figure 7A,B), DSI (Figure 7C-E) and normalized cocaine-induced psychomotor effects (Figure 7F), which is not distinguishable between unstressed and MAOA^*Neo*^ ES mice. Finally, indirect activation of CB1 receptors by pharmacologically enhancing 2-AG tone through in vivo MAGL inhibition (JZL 184: 8 mg/kg, i.p. 7 days, 30 min before each cocaine injection), restored synaptic properties and plasticity at inhibitory synapses on VTA dopamine cells MAOA^*Neo*^ ES mice (Figure 8A-E) as well as their locomotor responses to cocaine (Figure 8F).

**Figure 7.**
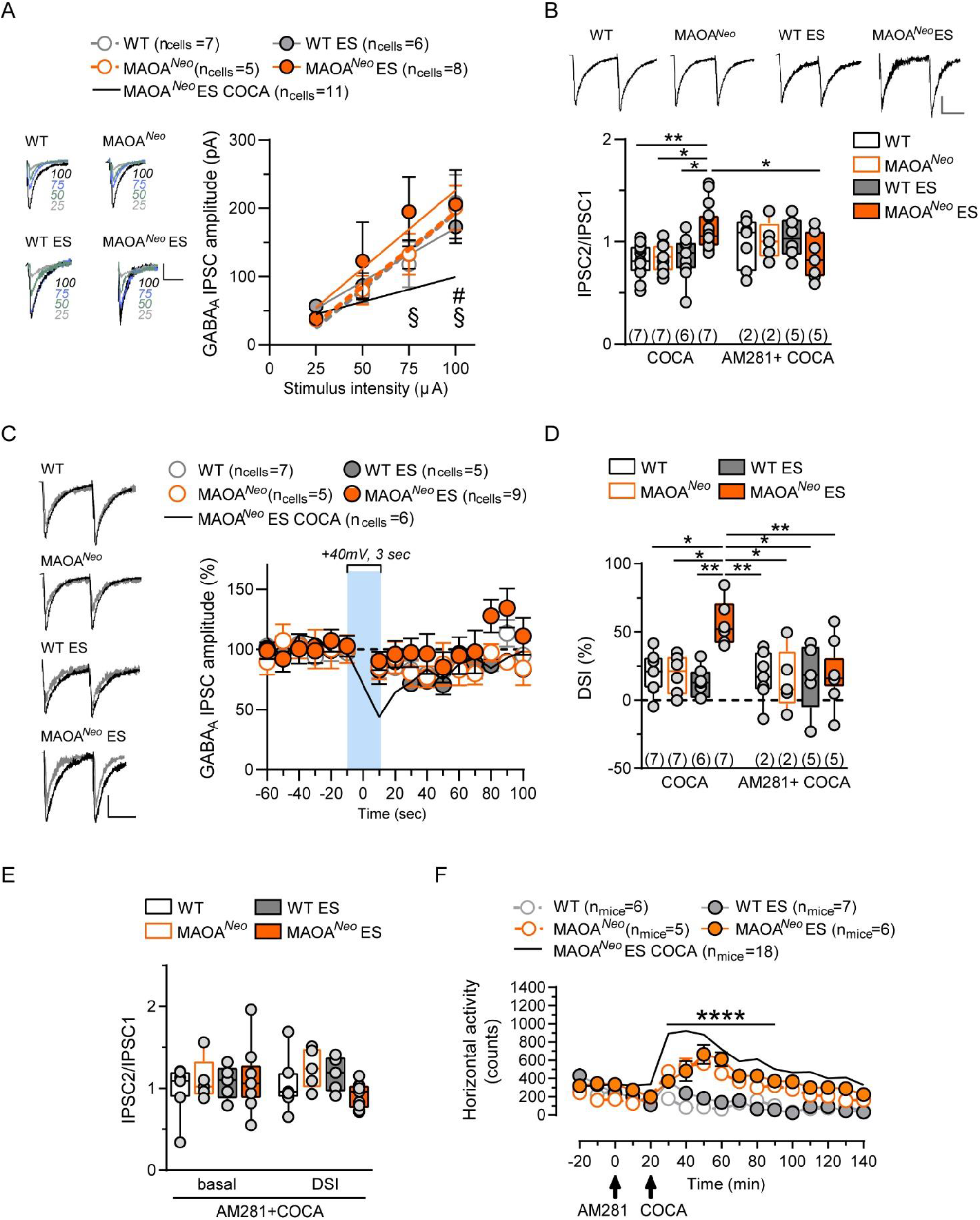
*In vivo* pretreatment with CB1R antagonist, AM281, restores synaptic properties and plasticity of inhibitory transmission on VTA DA neurons and normalizes cocaine-induced hyperlocomotion in MAOA^*Neo*^ mice subjected to ES. **(A)** Input–output relationships of GABAA-mediated IPSCs shows the effect of pretreatment with CB1R antagonist, AM281 (2.5 mg/kg, i.p) 20 min before each intraperitoneal cocaine injection (COCA, 15 mg/kg) on midbrain DA neurons 24 hrs after the last repeated treatment (1 week). AM281 abolishes cocaine-induced reduction in the amplitude of GABAA IPSC in MAOA^*Neo*^ ES mice and the differences among groups, WT (n_cells,mice_=7,2), MAOA^*Neo*^ (n_cells,mice_=5,2), WT ES (n_cells,mice_=6,5) and MAOA^*Neo*^ ES (n_cells,mice_=8,5) mice, (Two-way RM ANOVA, Stimulus intensity effect F_(3,96)_=34,99, p<0,0001, Gene x ES x Trt effect F_(4,32)_=1,44, p=0,24, Subject effect F_(32,96)_=4,74, p<0,0001, Interaction F_(12,96)_=1,75, p=0,067, Tukey’s multiple comparisons post hoc test, 75 μA, ^§^MAOA^*Neo*^ ES COCA vs MAOA^*Neo*^ ES AM281+COCA, p<0,05, 100 μA, ^#^MAOA^*Neo*^ ES COCA vs WT AM281+COCA, p<0,05, ^§^MAOA^*Neo*^ ES COCA vs MAOA^*Neo*^ ES AM281+COCA, p<0,05). Each symbol represents the averaged value (± SEM) obtained from different cells. Insets show representative traces of GABAA IPSC recorded at each stimulus intensity. Calibration bar: 25 ms, 50 pA. (**B**) Graph panels summarizing the effect of pretreatment with AM281 on averaged paired-pulse ratio (IPSC2/IPSC1) for DA cells recorded in WT (n_cells,mice_=7,2), MAOA^*Neo*^ (n_cells,mice_=5,2), WT ES (n_cells,mice_=6,5) and MAOA^*Neo*^ ES (n_cells,mice_=7,5) mice (Two way ANOVA, Gene x ES effect F_(3,68)_=0,63, p=0,595, Treatment effect F_(1,68)_=1,70, p=1,196, Interaction F_(3,68)_=5,99, p=0,001, Tukey’s multiple comparisons post hoc test, **MAOA^*Neo*^ ES COCA vs WT COCA, p=0,002, *MAOA^*Neo*^ ES COCA vs MAOA^*Neo*^ COCA, p=0,019, *MAOA^*Neo*^ ES COCA vs WT ES COCA, p=0,01, *MAOA^*Neo*^ ES COCA vs MAOA^*Neo*^ ES AM281+COCA, p=0,059). Inset shows representative traces of paired GABAA IPSCs recorded from VTA putative DA neurons. Calibration bar: 25 ms, 50 pA. (**C**) Graph panel shows the effect of pretreatment with AM281 on time course of DSI 24 hr after the last repeated AM281+COCA treatment. Application of a depolarizing pulse from -70 to +40 mV for 3 sec on VTA DA neurons prevents DSI of GABAA IPSCs evoked by stimulating rostral afferents in WT (n_cells,mice_=7,2), MAOA^*Neo*^ (n_cells,mice_=5,2), WT ES (n_cells,mice_=5,5) and MAOA^*Neo*^ ES mice (n_cells,mice_=9,5) (Two way RM ANOVA, Time effect F_(17,408)_=3,89, p<0,0001, Gene x ES effect F_(4,24)_=4,24, p=0,009, Subject effect F_(24,408)_=2,64, p<0,0001, Interaction F_(68,408)_=1,11, p=0,272). GABAA IPSCs amplitude was normalized to the averaged value (dotted line) before depolarization. Each symbol represents the averaged value (± SEM) obtained from different cells. Insets show representative traces of IPSC recorded before (black) and after DSI (grey). Calibration bar, 25 ms, 50 pA. (**D**) Averaged data for DSI induced by depolarizing pulse with a duration 3 sec are plotted (Two way ANOVA, Gene x ES effect F_(3,44)_=4,35, p=0,009, Treatment effect F_(1,44)_=3,64, p=0,063, Interaction F_(3,44)_=3,51, p=0,023, Tukey’s multiple comparisons post hoc test, *MAOA^*Neo*^ ES COCA vs WT COCA, p=0,017, *MAOA^*Neo*^ ES COCA vs MAOA^*Neo*^ COCA, p=0,021, **MAOA^*Neo*^ ES COCA vs WT ES COCA, p=0,003, **MAOA^*Neo*^ ES COCA vs WT AM281+COCA, p=0,009, *MAOA^*Neo*^ ES COCA vs MAOA^*Neo*^ AM281+COCA, p=0,012, *MAOA^*Neo*^ ES COCA vs WT ES AM281+COCA, p=0,022, **MAOA^*Neo*^ ES COCA vs MAOA^*Neo*^ ES AM281+COCA, p=0,008). (**E**) Bar graph summarizing the averaged paired-pulse ratio (IPSC2/IPSC1) of GABAA IPSCs for all cells recorded after pretreatment of AM281 before (basal) and after (DSI) depolarization pulse (Two way ANOVA, Gene x ES effect F_(3,44)_=1,015, p=0,395, Basal vs DSI effect F_(1,44)_=0,27, p=0,604, Interaction F_(3,44)_=1,105, p=0,375). (**F**) Pretreatment of mice with CB1R antagonist AM281 for 1 week abolishes the cocaine-induced hyperlocomotion in MAOA^*Neo*^ ES mice (Two-way RM ANOVA followed by Tukey’s multiple comparisons test, Time effect F_(16,592)_=10,28, p<0,0001, Gene x ES x treatment effect F_(4,37)_ =17,11, p<0,0001, Subject effect F_(37,592)_=8,238, p<0,0001, Interaction F_(64,592)_=3,857, p<0,0001, n mice=6 WT, 5 MAOA^*Neo*^, 7 WT ES, 6 MAOA^*Neo*^ ES, 18 MAOA^*Neo*^ ES COCA). Graph shows spontaneous locomotor (horizontal) activity measured as counts (bin=10 min) before and after drugs injection at the last day of repeated treatment in a novel open field arena. Data are represented as mean ± SEM. Unless otherwise indicated, graphs show box- and-whisker plots (including minima, maxima and median values, and lower and upper quartiles) with each circle representing a single cell recorded (numbers in the brackets represent number of the mice).

**Figure 8.**
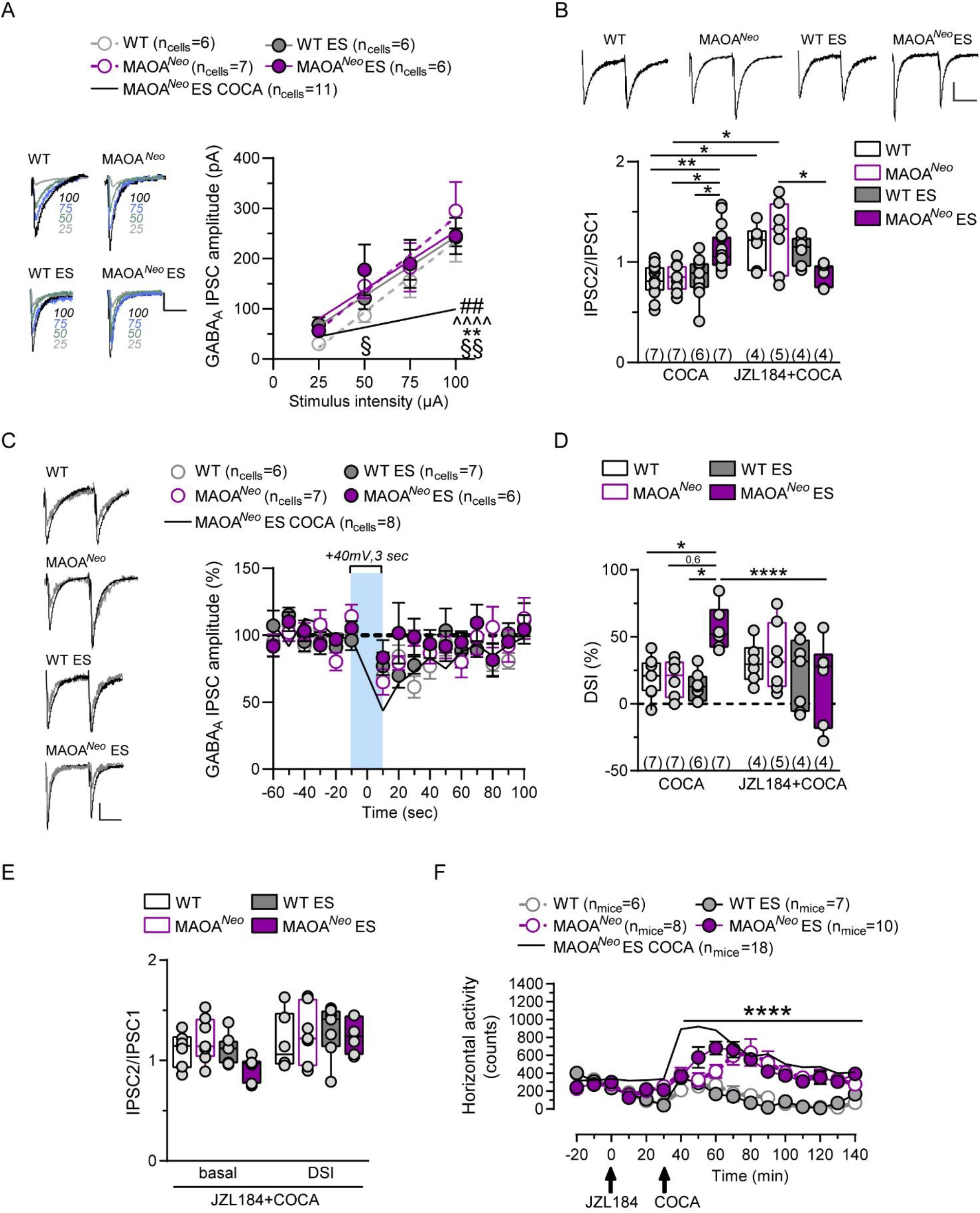
*In vivo* pretreatment with MAGL inhibitor, JZL184, restores synaptic properties and plasticity of inhibitory transmission on VTA DA neurons and reduces cocaine-induced hyperlocomotion in MAOA^*Neo*^ mice subjected to ES. (**A**) Input–output relationships of GABAA-mediated IPSCs shows the effect of pretreatment with MAG lipase inhibitor, JZL184, (8 mg/kg, i.p) 30 m before each intraperitoneal cocaine injection (COCA, 15 mg/kg) on midbrain DA neurons 24 hrs after the last repeated treatment (1 week). JZL184 abolishes cocaine-induced reduction in the amplitude of GABAA IPSC in MAOA^*Neo*^ ES mice and the differences among groups, WT (n_cells,mice_=8,4), MAOA^*Neo*^ (n_cells,mice_=7,5), WT ES and MAOA^*Neo*^ES (n_cells,mice_=6,4 per group) mice (Two-way RM ANOVA, Stimulus intensity effect F_(3,96)_=49,95, p<0,0001, Gene x ES x Trt effect F_(4,32)_=4,08, p=0,008, Subject effect F_(32,96)_=4,03, p<0,0001, Interaction F_(12,96)_=2,58, p=0,005, Tukey’s multiple comparisons post hoc test, 50 μA, ^§^MAOA^*Neo*^ ES COCA vs MAOA^*Neo*^ ES JZL184+COCA p≤0,05, 100 μA, ^##^MAOA^*Neo*^ ES COCA vs WT JZL184+COCA p<0,01, ^^^^MAOA^*Neo*^ ES COCA vs MAOA^*Neo*^ JZL184+COCA, p<0,0001, **MAOA^*Neo*^ ES COCA vs WT ES JZL184+COCA p<0,01, ^§§^MAOA^*Neo*^ ES COCA vs MAOA^*Neo*^ ES JZL184+COCA p<0,01). Each symbol represents the averaged value (± SEM) obtained from different cells. Insets show representative traces of GABAA IPSC recorded at each stimulus intensity. Calibration bar: 25 ms, 50 pA. **(B)** Graph panels summarizing the effect of pretreatment with JZL184 on averaged paired-pulse ratio (IPSC2/IPSC1) for DA cells recorded in WT (n_cells,mice_=8,4), MAOA^*Neo*^ (n_cells,mice_=7,5), WT ES and MAOA^*Neo*^ES (n_cells,mice_=6,4 per group) mice (Two way ANOVA, Gene x ES effect F_(3,69)_=0,41, p=0,75, Treatment effect F_(1,69)_=16,6, p=0,0001, Interaction F_(3,69)_=10,07, p<0,0001, Tukey’s multiple comparisons post hoc test, **MAOA^*Neo*^ ES COCA vs WT COCA, p=0,003, *MAOA^*Neo*^ ES COCA vs MAOA^*Neo*^ COCA, p=0,024, *MAOA^*Neo*^ ES COCA vs WT ES COCA, p=0,0135, *WT COCA vs WT JZL184+COCA, p=0,021, **WT COCA vs MAOA^*Neo*^ JZL184+COCA, p=0,0002, **MAOA^*Neo*^ COCA vs MAOA^*Neo*^ JZL184+COCA, p=0,001, ***WT ES COCA vs MAOA^*Neo*^ JZL184+COCA, p=0,0009, *MAOA^*Neo*^ JZL184+COCA vs MAOA^*Neo*^ ES JZL184+COCA, p=0,015). Inset shows representative traces of paired GABAA IPSCs recorded from VTA putative DA neurons. Calibration bar: 25 ms, 50 pA. (**C**) Graph panel shows the effect of pretreatment with JZL184 on time course of DSI on VTA DA neurons 24 hr after the last repeated JZL184+COCA treatment in WT (n_cells,mice_=6,4), MAOA^*Neo*^ (n_cells,mice_=7,5), WT ES (n_cells,mice_=7,4) and MAOA^*Neo*^ ES (n_cells,mice_=6,4) mice (Two way RM ANOVA, Time effect F_(15,405)_=6,75, p<0,0001, Gene x ES effect F_(4,27)_=1,031, p=0,409, Subject effect F_(27,405)_=2,189, p=0,0007, Interaction F_(60,405)_=1,04, p=0,396). GABAA IPSCs amplitude was normalized to the averaged value (dotted line) before depolarization. Each symbol represents the averaged value (± SEM) obtained from different cells. Panel shows representative traces of IPSCs recorded before and after DSI (grey). Calibration bar, 25 ms, 50 pA. (**D**) Averaged data for DSI induced by application of a depolarizing pulse from -70 to +40 mV for 3 sec on VTA DA neurons are plotted (Two way ANOVA, Gene x ES effect F_(3,44)_=1,69, p=0,183, Treatment effect F_(1,44)_=0,013, p=0,91, Interaction F_(3,44)_=4,78, p=0,005, Tukey’s multiple comparisons post hoc test, *MAOA^*Neo*^ ES COCA vs WT COCA, p=0,052, *MAOA^*Neo*^ ES COCA vs WT ES COCA, p=0,012, MAOA^*Neo*^ ES COCA vs MAOA^*Neo*^ COCA, p=0,061 ns, *MAOA^*Neo*^ES COCA vs MAOA^*Neo*^ES JZL184+COCA, p=0,038). (**E**) Bar graph summarizing the averaged paired-pulse ratio (IPSC2/IPSC1) of GABAA IPSCs for all cells recorded after pretreatment with JZL184, before (basal) and after (DSI) depolarization pulse (Two way ANOVA, Gene x ES effect F_(3,44)_=1,17, p=0,33, Basal vs DSI effect F_(1,44)_=6,86, p=0,012, Interaction F_(3,44)_=1,011, p=0,396). (**F**) Pretreatment of mice with MAGL inhibitor, JZL184, for 1 week abolishes the cocaine-induced hyperlocomotion in MAOA^*Neo*^ mice exposed to ES (Two-way RM ANOVA followed by Tukey’s multiple comparisons test, Time effect F_(4.146, 182.4)_=14,18, p<0,0001, Gene x ES x treatment effect F_(4, 44)_ =20,28, p<0,0001, Subject effect F_(44, 528)_=4,23, p<0,0001, Interaction F_(48, 528)_=5,2, p<0,0001, n mice=6 WT, 8 MAOA^*Neo*^, 7 WT ES, 10 MAOA^*Neo*^ ES, 18 MAOA^*Neo*^ ES COCA). Graph shows spontaneous locomotor (horizontal) activity measured as counts (bin=10 min) before and after drugs injection at the last day of repeated treatment in a novel open field arena. Data are represented as mean ± SEM. Unless otherwise indicated, graphs show box- and-whisker plots (including minima, maxima and median values, and lower and upper quartiles) with each circle representing a single cell recorded (numbers in the brackets represent number of the mice).

Collectively, these results suggest that in MAOA^*Neo*^ ES mice repeatedly treated with cocaine, DAD2 and CB1 receptors cooperate to diminish the strength of GABA transmission on VTA dopamine neurons and to enhance sensitivity to cocaine-induced hyperlocomotion.

## Discussion

In the present study, we provide evidence that repeated in vivo exposure to cocaine produces a long lasting increase in its psychomotor stimulant effects in individuals whose susceptibility is driven by both genetic (low activity of MAOA) and environmental (early life adversity) influences. The enhanced responsiveness to psychostimulant actions of cocaine exhibited by pre-adolescent MAOA^*Neo*^ ES mice is associated with long lasting adaptations specifically at inhibitory afferents on dopamine neurons of the VTA, an important locus of cocaine actions. These persistent functional modifications at GABAergic synapses on dopamine cells require the interplay between dopamine and the endocannabinoid 2-AG, which is key for the cross-sensitization of cocaine with environmental (ES) and genetic (MAOA^*Neo*^) risk factors.

These findings support and extend previous studies showing that early life adversity per se can enhance sensitization to cocaine [(Vannan et al., 2018) and references therein], although this effect is often moderated by genetic factors (Fernandez-Castillo et al., 2022). Exposure to early life stress is a critical environmental risk factor for the development of drug-related issues (Enoch, 2011; Vannan et al., 2018; Fite et al., 2019b; Alves et al., 2020). In rodents, the use of maternal separation as a model of early life adversity has provided important information about the mechanisms underlying susceptibility to drug misuse later in life (Delavari et al., 2016; Viola et al., 2016; Vannan et al., 2018; Ganguly et al., 2019; Castro-Zavala et al., 2020; Castro-Zavala et al., 2021). A dysregulation of dopamine signaling in mesolimbic pathway has been associated to cross-sensitization between maternal separation and cocaine exposure (Brake et al., 2004; Enoch, 2011; Castro-Zavala et al., 2021). Accordingly, marked and lasting molecular alterations affect the responsiveness of mesolimbic dopamine neurons to subsequent exposure to stress and psychostimulants and might predispose to the progression from cocaine use to abuse and dependence. This is particularly relevant when it occurs during vulnerable periods of life such as adolescence (Brake et al., 2004; Delavari et al., 2016; Viola et al., 2016; Vannan et al., 2018; Ganguly et al., 2019; Castro-Zavala et al., 2020; Castro-Zavala et al., 2021).

Our data substantiate the notion that not all the individuals who experience early life stress will develop mental illness, particularly substance use disorders, and highlight the role of other mediating factors in worsening one’s lifetime trajectory, such as G x E interactions (Vink, 2016). Accordingly, among the several genes implicated in the susceptibility to substance use disorders, those related to monoamines (i.e., dopamine, serotonin, and norepinephrine) have been shown to serve important roles (Volkow et al., 2007; Howell and Kimmel, 2008), particularly when associated to early life adversity (van Amsterdam et al., 2006; Lea and Chambers, 2007). Hence, clinical studies show that the interplay between low activity of MAOA gene and lifetime stress enhances the risk for early experimentation with drugs of abuse (Vanyukov et al., 2004; Vanyukov et al., 2007; Stogner and CL, 2013; Fite et al., 2019a; Fite et al., 2019b), besides the one for aggressiveness (Caspi et al., 2002; Viding and Frith, 2006; Guo et al., 2008; Weder et al., 2009; Checknita et al., 2015). Here, we propose that MAOA hypomorphic transgenic mice exposed to an early life stress (MAOA^*Neo*^ ES mice) also display a vulnerability for developing cocaine use disorders, in line with the “plasticity alleles” hypothesis (Belsky and Beaver, 2011). We, therefore, extend the overall risk of developing psychopathology (i.e., aggressive behavior) (Frau et al., 2019a; Godar et al., 2019) exhibited by this mouse model. This has a unique translational advantage to unveil the underpinnings of the actions of such a heavily-abused substance in an at-risk segment of population where the interplay of this gene with lifetime stress increases the predisposition to progress into a substance use disorder (Vanyukov et al., 2004; Vanyukov et al., 2007; Stogner and CL, 2013; Fite et al., 2019a; Fite et al., 2019b). Notably, this mouse model recapitulates a complex G x E interaction, which produces long lasting brain adaptations in monoaminergic systems (Frau et al., 2019a; Godar et al., 2019), heavily implicated in cocaine’s rewarding actions. Accordingly, VTA dopamine neurons from MAOA^*Neo*^ ES mice display an increased responsiveness to excitatory stimuli (Frau et al., 2019a), a key neural adaptation contributing to the effects of stress and to core features of drug addiction (Saal et al., 2003; Bellone et al., 2021), thereby imprinting a susceptibility trait.

The observation that repeated in vivo cocaine exposure does not enhance the strength at excitatory synapses on VTA dopamine neurons in MAOA^*Neo*^ ES mice suggests that this G x E interaction disrupts the normal, presumably adaptive, relationship between locomotor response to repeated cocaine exposure and increased synaptic strength in the VTA. Whether or not this depends on a saturated potentiation of excitatory inputs onto dopamine neurons, similarly to that observed following in vivo exposure to cocaine itself (Ungless et al., 2001; Borgland et al., 2004), remains to be established. Nonetheless, not only we show that the enhancement of synaptic strength at excitatory synapses in the VTA is not required for the expression of cocaine behavioral sensitization in MAOA^*Neo*^ ES mice, but also that a long lasting decrease in the efficacy of GABAergic transmission on dopamine neurons plays a pivotal role. Accordingly, cocaine is more potent and more effective in decreasing the amplitude of GABAA IPSCs in MAOA^*Neo*^ ES mice. In rats, repeated in vivo exposure to cocaine reduces GABA synaptic efficacy (Liu et al., 2005; Pan et al., 2008b), which increases the chance of eliciting an action potential in VTA dopamine neurons and facilitates the induction of long-term potentiation at excitatory synapses (Liu et al., 2005). Conversely, our results from MAOA^*Neo*^ ES mice also substantially extend the observation of a cooperative interaction between endocannabinoids and dopamine in the regulation of GABA transmission on rat VTA dopamine neurons only upon repeated cocaine exposure in vivo (Pan et al., 2008b) via activation of dopamine D2 (DAD2) and CB1 receptors (Pan et al., 2008a, b). Hence, we show that in MAOA^*Neo*^ ES mice concurrent in vivo cocaine administration with either DAD2 or CB1 antagonist prevents cocaine-induced i) reduction of the amplitude of maximal GABAA IPSCs, ii) decreased probability of GABA release, iii) expression of DSI, and iv) hyperlocomotion. Similarly, in MAOA^*Neo*^ ES mice, concurrent in vivo cocaine treatment with the MAGL inhibitor JZL184, by enhancing 2-AG levels and thereby acting as a CB1 receptor indirect agonist, averts the effects induced by repeated cocaine exposure on GABA synaptic efficacy, DSI and locomotor activity. Collectively, these data indicate that behavioral sensitization and cocaine-induced reduction of GABAergic synaptic efficacy and DSI in VTA dopamine cells share common mechanisms and molecular targets for both cocaine and the endogenous retrograde signal, which involve the activation of DAD2 and CB1 receptors.

Remarkably, repeated in vivo cocaine exposure enables a form of 2-AG-mediated short term synaptic plasticity at these synapses (i.e., DSI) that is expressed by VTA dopamine neurons only in MAOA^*Neo*^ ES mice, a phenomenon that persists one week after the last cocaine administration. Our findings confirm that repeated in vivo cocaine administration alone could not empower the expression of DSI in rat VTA dopamine cells (Pan et al., 2008b). In MAOA^*Neo*^ ES mice, DSI cannot be ascribed to a larger number of presynaptic CB1 receptors, or a greater effect produced upon their activation. Conversely, the observation that 2-AG phasic signal is enhanced in either time and/or space at these inhibitory afferents impinging upon VTA dopamine neurons is consistent with previous reports showing that the strength of retrograde synaptic depression is determined by 2-AG rate of degradation (Pan et al., 2009; Yoshida et al., 2011; Melis et al., 2013; Melis et al., 2014). Particularly, pharmacological blockade of MAGL resulted in a DSI similarly expressed by dopamine cells of all experimental groups. Since MAOA^*Neo*^ ES mouse dopamine neurons per se do not express DSI (data not shown), it is important to elucidate how cocaine, when repeatedly administered, affects sensitivity and/or activity and/or expression of MAGL exclusively in MAOA^*Neo*^ ES mice. On one hand, our results are in contrast with previous reports showing that, in C57Bl6/J mice, repeated cocaine sensitization decreases 2-AG synthesis/degradation ratio, thus suggesting a reduced 2-AG production that is associated with a compensatory upregulation of CB1 receptors in discrete brain regions (Blanco et al., 2014; Palomino et al., 2014; Blanco et al., 2016). However, one must point out that genetic mouse background, cocaine dose and duration of regimen, and brain regions examined do differ from ours and this might help resolving these discrepancies. Particularly, 129S6 mice are resistant to psychostimulant effects of cocaine (Thomsen and Caine, 2011). Nonetheless, it worth noting that a similar enhancement in phasic 2-AG signaling mediating DSI expressed by rat VTA dopamine neurons is associated to innate alcohol preference (Melis et al., 2014) as well as to a faster acquisition in cannabinoid-self administration (Melis et al., 2013). Remarkably, not only an important consequence of 2-AG-mediated DSI is to transiently silence inhibitory inputs onto dopamine cells, but also to increase their responsiveness to excitatory stimuli, thereby priming them for subsequent plasticity induction. One, therefore, is tempted to speculate that this 2-AG-mediated plasticity might contribute to vulnerability traits to some behavioral actions of different drugs of abuse.

Several limitations of the present study should be acknowledged. First, our behavioral analysis for the predisposition to cocaine abuse is curtailed to cocaine-induced hyperlocomotion. Second, our recordings were carried out from putative dopamine neurons within lateral portion of the VTA, which largely projects to the lateral shell of the Nucleus Accumbens (Lammel et al., 2011), so it is likely that these putative dopamine neurons would mainly project to this region. Third, besides blocking membrane transporters for dopamine, cocaine it also blocks the uptake of serotonin and norepinephrine, as well as ligand- and voltage-gated channels (Uhl et al., 2002). Therefore, we cannot rule out the contribution of other monoaminergic systems and ion-channels in both synaptic and behavioral outcomes of MAOA^*Neo*^ ES mice. These limitations notwithstanding, our results gain insights into the mechanisms influencing the overall risk for cocaine use disorder, and identify molecular targets (i.e., DAD2, CB1, MAGL) that could prove effective in age-specific management of such disorder.

In conclusion, our findings provide evidence that multifactorial conditions driven by specific genetic (low activity of MAOA), biological (age) and environmental (early life stress, cocaine) influences can produce profound and long lasting molecular alterations leading to a reduced GABA input to VTA putative dopamine cells, and an enhanced 2-AG-mediated transmission at these synapses that have important behavioral associations, including vulnerability to cocaine addiction.

## Materials and Methods

### Animals

Male MAOA^*Neo*^ mice were generated from 129S6 genetic background by mating primiparous MAOA^*Neo*^ heterozygous females with wild-type (WT) sires, as previously described (Bortolato et al., 2011). Since Maoa is an X-linked gene, male offspring of MAOA^*Neo*^ mice dams were either MAOA^*Neo*^ or WT. Pregnant MAOA^*Neo*^ mice were single housed 3 days prior to parturition. Only litters with more than 4 pups (and at least 2 males) were used, and all litters with more than 8 pups were culled to 8 at postnatal day (PND) 1 to assure uniformity of litter size. Litter effects were minimized by using mice from at least six different litters for behavioral and electrophysiological experiment. Bedding was changed in all cages at PND 7 and PND 14, and mice were weaned at PND 21. Animals were grouped-housed in cages with food and water available ab libitum, except during the periods of social isolation, in which they were isolated in their home cage with ab libitum access to food and water. The room was maintained at 22°C, on a 12 hr/12hr light/dark cycle from 7 a.m. to 7 p.m. Behavioral and electrophysiological experiments occurred between 10 a.m and 6 p.m. during the light phase of the light/dark cycle.

All procedures were performed in accordance with the European legislation (EU Directive, 2010/63) and were approved by the Animal Ethics Committees of the University of Cagliari and by Italian Ministry of Health (auth. n. 621/2016-PR). All efforts were made to minimize animal pain and discomfort and to reduce the number of animals used. Male MAOA^*Neo*^ mice on 129S6 genetic background and their WT counterparts were used in this study.

### Early stress procedures (ES)

To simulate child neglect and maltreatment, male pups were subjected to a daily stress regimen of maternal separation and saline injection respectively, during the first postnatal week (from PND 1 to 7; Figure 1A), as previously described (Godar et al., 2019), hereafter designated as early life stress (ES). During maternal separation, pups were removed from the home cage and placed into a new cage in a separate temperature-controlled (25 °C) room for 2–3 hr/day in a pseudorandom fashion and at different times during the light cycle. Physiological saline injections were performed using a micro-injector connected to a Hamilton syringe (10 μL/g body weight). The stressfulness of intraperitoneal injection was ensured by verifying the response from each pup, which typically consisted of rapid limb and head movements after the puncture. At PND 22, mice were singly housed for 5 days before performing behavioral and electrophysiological experiments (Figure 1A). This regimen was implemented to enhance the translational relevance of the animal model (Godar et al., 2019).

### Open field test

Locomotor activity was tested in the open field (OF), consisting of a clear Plexiglas square arena (11 × 11 × 30 cm) covered by clear Plexiglas plate, as previously described (Frau et al., 2019b). Each box had a set of photocells located at right angles to each other and projecting horizontal infrared beam. Mice were placed into the central arena allowed to freely explore the OF. Basal locomotor activity was measured as total number of sequential infrared beam breaks in the horizontal sensor, recorded every 5 minutes (min). To evaluate the effect of cocaine on locomotor behavior, animals were first habituated to the apparatus for 15-30 min to obtain a stable measure of baseline activity, then after cocaine administration the locomotor activity was assessed for 120 min. We recorded the response of mice in locomotor activity (a) after a single cocaine administration, (b) the last day of repeated cocaine exposure (7 days) and (c) after a challenge with cocaine following 1 week of withdrawal from cocaine repeated exposure (Figure 1A).

### Ex vivo electrophysiology

The preparation of VTA slices was as described previously (Frau et al., 2019a). Briefly, horizontal midbrain slices (250 μm) were prepared 24 hr after repeated cocaine in vivo exposure and one week after the last cocaine injection, respectively, from MAOA^*Neo*^ and WT mice (PND 28-30). Mice were anesthetized with isoflurane until loss of righting reflex. VTA slice were prepared in ice cold (4–6°C) low-Ca2+ solution containing the following (in mM): 126 NaCl, 1.2 KCl, 1.2 NaH2PO4, 1.2 MgCl2, 0.625 CaCl2, 18 NaHCO3 and 11 glucose (304-306 mOsm). Immediately after cutting, slices were allowed to recover for at least 1 hr at 36–37 °C until recording and superfused with artificial cerebrospinal fluid (aCSF) containing (in mM): 126 NaCl, 1.6 KCl, 1.2 NaH2PO4, 1.2 MgCl2, 2.4 CaCl2, 18 NaHCO3 and 11 glucose (304-306 mOsm). All solutions were saturated with 95% O2 and 5% CO2. Cells were visualized with an upright microscope with infrared illumination (Axioskop FS 2 plus; Zeiss), and whole-cell patch-clamp recordings were made by using an Axopatch 200 B amplifier (Molecular Devices). Voltage-clamp recordings of evoked excitatory post-synaptic currents (EPSCs) were made with electrodes filled with a solution containing the following (in mM): 117 Cs methanesulfonic acid, 20 HEPES, 0.4 EGTA, 2.8 NaCl, 5 TEA-Cl, 2.5 Mg2ATP, and 0.25 Mg2GTP, pH 7.2–7.4, 275–285 mOsm. Picrotoxin (100 μm) was added to the aCSF to block GABAA receptor-mediated inhibitory post-synaptic currents (IPSCs). Voltage-clamp recordings of evoked IPSCs were made with electrodes filled with a solution containing the following (in mM): 144 KCl, 10 HEPES, 3.45 BAPTA, 1 CaCl2, 2.5 Mg2ATP and 0.25 Mg2GTP, pH 7.2–7.4, 275–285 mOsm. All GABAA IPSCs were recorded in the presence of 6-cyano-2,3-dihydroxy-7-nitro-quinoxaline (10 μM) and 2-amino-5-phosphonopentanoic acid (AP5; 100 μM) to block AMPA and NMDA -receptors-mediated synaptic currents, respectively. Experiments were begun only after series resistance had stabilized (typically 15–40 MΩ). Series and input resistance were monitored continuously on-line with a 5 mV depolarizing step (25 ms). Data were filtered at 2 kHz, digitized at 10 kHz, and collected on-line with acquisition software (pClamp 10; Molecular Devices). Dopamine neurons from the lateral portion of the posterior VTA were identified by the following criteria: cell morphology and anatomical location to the medial terminal nucleus of the accessory optic tract, the presence of a large hyperpolarization activated current (Ih ≥ 100 pA) assayed immediately after break-in using 13 incremental 10 mV hyperpolarizing steps from a holding potential of -70 mV (Frau et al., 2019b). A bipolar, stainless steel stimulating electrode (FHC) was placed ∼100 – 200 μm rostral to the recording electrode and was used to stimulate at a frequency of 0.1 Hz. Paired stimuli were given with an interstimulus interval of 50 ms and the ratio between the second and the first postsynaptic currents (PSC2/PSC1) was calculated and averaged for a 5 min baseline (Melis et al., 2002). NMDA EPSCs were evoked while holding cells at +40 mV. The AMPA EPSC was isolated after bath application of the NMDA antagonist D-AP5 (100 μM). The NMDA EPSC was obtained by digital subtraction of the AMPA EPSCs from the dual (AMPA + NMDA-mediated) EPSC (Ungless et al., 2001). The depolarizing pulse used to evoke depolarization-induced suppression of inhibition (DSI) was a 3 s step to +40 mV from holding potential (Melis et al., 2009). This protocol was chosen on the evidence of a manifest eCB tone when VTA DA cells are held at + 40 mV for 3 sec (Melis et al., 2004; Melis et al., 2009). The magnitude of DSI was measured as a percentage of the mean amplitude of consecutive IPSCs after depolarization (acquired between 5 and 15 sec after the end of the pulse) relative to that of 6 IPSCs before the depolarization. Bath application of WIN55,212-2 (WIN) was performed as follows: WIN was applied for 5 min at the lowest concentration, then, another 5 min with the next increasing WIN concentration. The effect of WIN on GABAA IPSCs was taken at the 5th min of bath application and normalized to the baseline (5 min before drug application). We chose this protocol because it has been shown that at physiological temperatures WIN-induced effects on GABAA IPSCs recorded from VTA DA neurons reached their peak at this time. The effect of JZL184 (100 nM) on GABAA IPSCs was taken after 5 min bath application (Melis et al., 2013). Each hemi-slice received only a single drug exposure. All drugs were dissolved in DMSO when it was needed. The final concentration of DMSO was < 0.01 %.

### Drug treatment

Cocaine hydrochloride was purchased from Johnson Matthey, sulpiride was purchased from Teofarma, AM281 and JZL184 were purchased from Tocris Bioscience, WIN55,212-2 was purchased from Sigma-Aldrich. Cocaine hydrochloride (15 mg/kg) and sulpiride (10 mg/kg) were dissolved in 0.9% saline solution, AM281 (2.5 mg/kg) was dissolved in a dimethyl sulfoxide (DMSO) and saline solution (1:3 v/v). JZL184 (8 mg/kg) was prepared as a saline:ethanol:cremophor (18:1:1 v/v/v) solution. Sulpiride and AM281 were administered 20 min before cocaine injection, JZL184 was administered 30 min before cocaine injection. All drugs were administered intraperitoneally (i.p.) in a volume of 10 ml/kg.

### Statistical analysis

All the numerical data are given as mean ± SEM. Statistical analysis was performed using GraphPad Prism (version 6.01). Statistical outliers were identified using Grubb’s test (α = 0.05) and excluded from analyses. Data were compared and analyzed utilizing Student’s t-test, two-way ANOVA, or two-way RM ANOVA, when appropriate, followed by Tukey’s or Bonferroni’s post hoc test, as specified in each figure. Significance threshold was set at 95% confidence interval.

## Acknowledgements

The authors thank R. Frau, K. McPherson and P. Devoto for discussions and comments on the manuscript, and M. Tuveri, S. Aramo, and B. Tuveri for their skillful assistance. The present study was supported by the the Region of Sardinia (RASSR32909 to M.M.) and the University of Cagliari (RICCAR 2017 and 2018 to M.M.)

